## Supplementary Figure 1 for "Safety and data quality of EEG recorded simultaneously with multi-band fMRI"

### Supplementary Materials

#### ***Comparison of functional activations in a localizer task between the MB sequence and the SB-COV sequence.***

We validated that our MB protocol had sufficient functional contrast-to-noise ratio, by comparing it against a SB sequence optimized for brain coverage, which we name SB-COV. Considering the robust prior evidence [1,2], the current demonstration in a single subject should be considered only a confirmatory proof of principle.

The SB-COV sequence used TR: 1800 ms, TE: 25 ms, flip angle: 90, 34 slices, slice thickness 3 mm, fat saturation on, GRAPPA acceleration factor: 2, bandwidth: 2470 Hz per pixel, and resolution: 92 x 92, FoV: 230x230mm. A single pilot subject ran through a functional localizer task designed to identify auditory-, face-, and object-processing areas for each imaging sequence. The localizer consisted of two tasks, first a passive listening task to target auditory areas [3], followed by a 1-back task using randomized blocks of Ekman faces [4] and household objects to target face-specific [5] and object-specific [6] areas, respectively (15 trials per block, 9 emotion blocks equally split between sad/angry/neutral expressions, and 3 object blocks).

FMRI data was preprocessed and analyzed in MATLAB 2018b using SPM12 (revision 7487) [7]. Preprocessing included realignment, co-registration, normalization to MNI stereotaxic space [8], and smoothing (FWHM=6mm). The contemporary SB-COV sequence additionally underwent slice time correction prior to realignment. Serial autocorrelations were corrected using a FAST model for the MB sequence and an autoregressive AR(1) model for the SB-COV sequence. The general linear model included one regressor per stimulus type and six rigid-body head motion parameters. As expected in accord with prior evidence, for all three contrasts, the MB sequence demonstrated increased functional contrast-to-noise ratio (Fig S1).

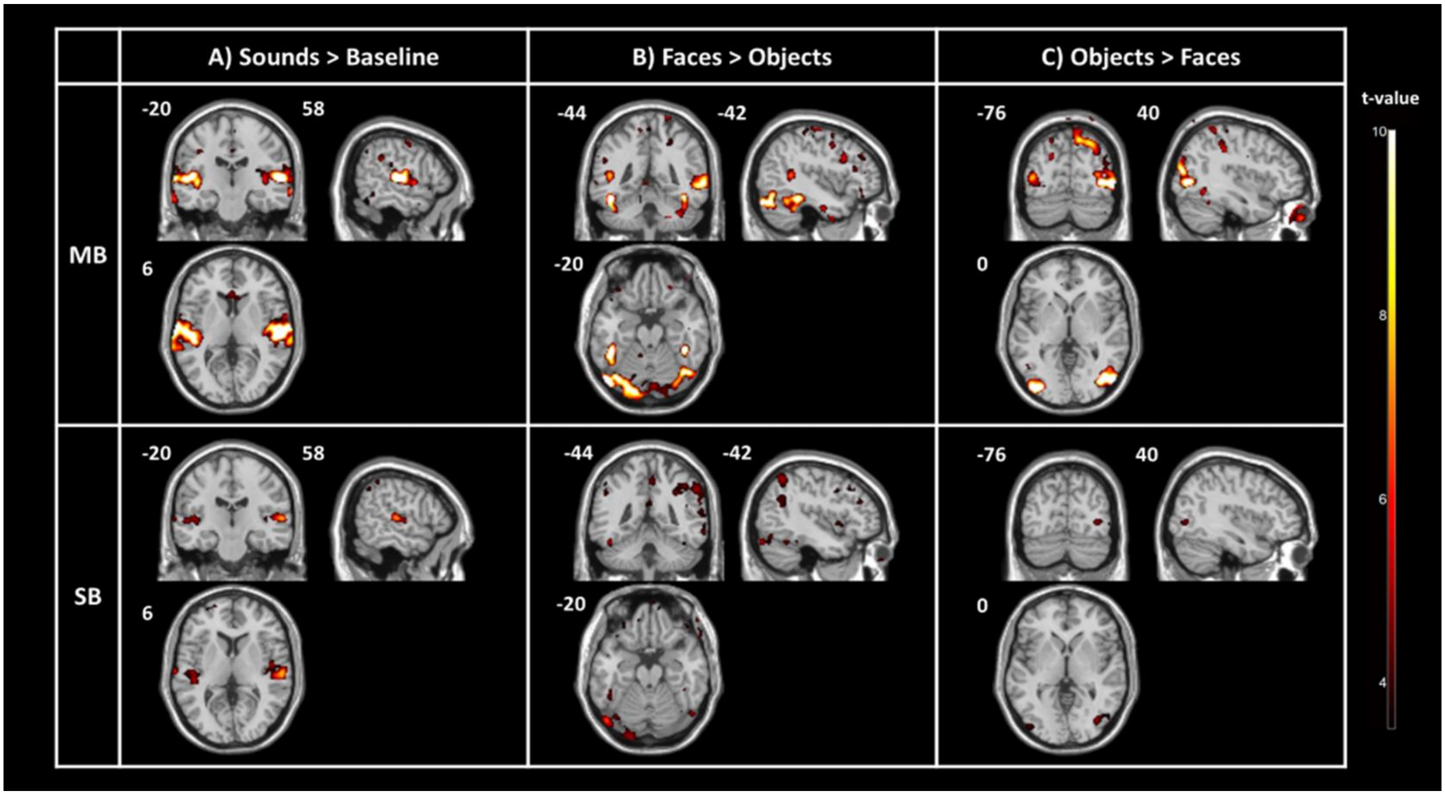


***Fig S1.*** *The visualized contrasts correspond to sounds>implicit baseline (A), faces>objects (B), and objects>faces (C) (threshold for visualization purposes: voxel-wise p<0.001 uncorrected). For each contrast, top and bottom rows (MB and SB-COV sequence, respectively) show equivalent slices for comparison. Activations comparable or stronger for the MB sequence are observed at the auditory cortex (A), the fusiform face area (B), and lateral occipital object-specific areas (C).*

**Supplementary Figure References**

1. Demetriou L, Kowalczyk OS, Tyson G, Bello T, Newbould RD, Wall MB. A comprehensive evaluation of increasing temporal resolution with multiband-accelerated protocols and effects on statistical outcome measures in fMRI. NeuroImage. 2018;176: 404–416. doi:10.1016/j.neuroimage.2018.05.011

2. Todd N, Moeller S, Auerbach EJ, Yacoub E, Flandin G, Weiskopf N. Evaluation of 2D multiband EPI imaging for high-resolution, whole-brain, task-based fMRI studies at 3T: Sensitivity and slice leakage artifacts. NeuroImage. 2016;124: 32–42. doi:10.1016/j.neuroimage.2015.08.056

3. Sadaghiani S, Hesselmann G, Kleinschmidt A. Distributed and Antagonistic Contributions of Ongoing Activity Fluctuations to Auditory Stimulus Detection. Journal of Neuroscience. 2009;29: 13410–13417. doi:10.1523/JNEUROSCI.2592-09.2009

4. Ekman P, Friesen W. Pictures of Facial Affect. Palo Alto, CA: Consulting Psychologists Press; 1976.

5. Berman MG, Park J, Gonzalez R, Polk TA, Gehrke A, Knaffla S, et al. Evaluating Functional Localizers: The Case of the FFA. Neuroimage. 2010;50: 56–71. doi:10.1016/j.neuroimage.2009.12.024

6. Grill-Spector K, Kourtzi Z, Kanwisher N. The lateral occipital complex and its role in object recognition. Vision Research. 2001;41: 1409–1422. doi:10.1016/S0042-6989(01)00073-6

7. Penny W, Friston K, Ashburner J, Kiebel S, Nichols T. Statistical Parametric Mapping: The Analysis of Functional Brain Images. 1st ed. Cambridge, MA: Academic Press; 2006.

8. Mazziotta JC, Toga AW, Evans A, Fox P, Lancaster J. A probabilistic atlas of the human brain: theory and rationale for its development. The International Consortium for Brain Mapping (ICBM). Neuroimage. 1995;2: 89–101. doi:10.1006/nimg.1995.1012
